# Neuropathic pain has sex-specific effects on oxycodone seeking and drug-seeking ensembles in the dorsomedial prefrontal cortex

**DOI:** 10.1101/2023.07.06.548045

**Authors:** Bailey C. Sarka, Shuai Liu, Anjishnu Banerjee, Cheryl L. Stucky, Qing-song Liu, Christopher M. Olsen

**Affiliations:** Department of Pharmacology and Toxicology, Medical College of Wisconsin; Neuroscience Research Center, Medical College of Wisconsin; Division of Biostatistics, Medical College of Wisconsin; Department of Cell Biology, Neurobiology and Anatomy, Medical College of Wisconsin

**Keywords:** opioid, self-administration, drug seeking, neuropathic pain, electrophysiology

## Abstract

Approximately 50 million Americans suffer from chronic pain, and opioids are commonly prescribed for such individuals. Unfortunately, nearly a quarter of chronic pain patients have reported misusing their prescription. We are investigating the effect of chronic pain on drug-seeking behavior at the neuronal level. Repeated drug-seeking is associated with reactivation of an ensemble of neurons sparsely scattered throughout the dorsomedial prefrontal cortex (dmPFC). Prior research has demonstrated that chronic pain increases intrinsic excitability of dmPFC neurons, which may increase the likelihood of reactivation during drug seeking. We tested the hypothesis that chronic pain would increase oxycodone seeking behavior, and that the pain state would differentially increase intrinsic excitability in dmPFC drug seeking ensemble neurons. TetTag mice self-administered intravenous oxycodone. After 7 days of forced abstinence, a drug seeking session (extinction conditions) was performed and the ensemble was tagged. Mice received spared nerve injury (SNI) to induce chronic pain during the period between a first and second seeking session, and we measured persistence of seeking between the two sessions to determine if the SNI exacerbated seeking. Following the second seeking session we performed electrophysiology on individual neurons within the dmPFC to assess intrinsic excitability of the drug-seeking ensemble and non-ensemble neurons. We found significant sex differences in the effect of SNI on oxycodone seeking and electrophysiology, such that the induction of chronic pain could modulate seeking behavior in mice that have previously self-administered oxycodone prior to injury.

**Highlights:**

- Oxycodone seeking was higher in females following SNI that came *after* the 10-day SA timeline.
- An increase in intrinsic excitability was detected among non-ensemble neurons from female mice that received SNI, and this correlated with an increase in seeking behavior.

## INTRODUCTION

The opioid epidemic reached new heights in 2020 when the CDC reported that nearly 70,000 lives were lost due to a drug overdose involving an opioid, making up 75% of all overdoses accounted for (1). The pandemic exacerbated an already existing problem for individuals misusing opioids (2,3). Some populations of people remain particularly vulnerable to opioid misuse, including those with chronic pain (4). Nearly 50 million people worldwide struggle with chronic pain (5), and opioids are commonly prescribed to help patients cope with their discomfort (6). Chronic pain can manifest in the wake of an acute injury and is marked by persistent existing pain (even beyond tissue repair at site of injury) and is accompanied by allodynia and hyperalgesia (7). Because chronic pain is persistent, prescriptions may extend beyond a year, and many patients require dose increases to maintain analgesic effects as tolerance builds up (6). It has been reported that approximately 21-29% of chronic pain patients misuse their prescription opioids (8).

Drug-seeking behavior can be initiated by “triggers.” These include stress (9), people that are associated with one’s drug use, objects, or the environment (10-12). These triggers can lead to the behavioral sequence to obtain the drug because there is a learned association between the administration of the drug, the physiological response, and the environment in which the drug is taken (6,13). The rodent dorsomedial prefrontal cortex (dmPFC) responds to these triggers to initiate drug seeking behavioral sequences (13). The dmPFC is also engaged during the transition from acute to chronic pain and is thought to contribute to the emotional effects of chronic pain (14,15). Taken together, it becomes evident that there is an overlap in this brain region regarding drug-seeking behavior and the affective experience of pain, although there is a lack of studies examining this overlap at the cellular level.

Within the dmPFC, there is evidence that a specific population of neurons, or ensemble, controls persistent substance seeking (16-19). Electrophysiological measures of pyramidal neurons in the prelimbic (PrL) region have shown that the induction of neuropathic pain in rats leads to an increase in intrinsic excitability (20). This change could be expected to increase responsiveness of dmPFC neurons to afferent inputs, such as those from drug-associated triggers, but it is unknown if these changes are happening to drug seeking ensemble neurons and if the induction of a pain state can increase drug seeking. Prior work has demonstrated that chronic pain (and inflammatory pain (21)) can elevate opioid seeking after self-administration, but these studies induced the pain state prior to opioid self-administration (22). In this design, opioids may be acting through both positive and negative (analgesic) reinforcement mechanisms. These studies concluded that the addition of negative reinforcement mechanisms was responsible for the chronic pain-associated increase in opioid seeking, although this hypothesis has not been tested. Here, we sought to examine the effects of chronic neuropathic pain on opioid seeking when the pain state occurred following opioid self-administration. We have also included females in this study, considering that there is a higher frequency of females and/or higher pain ratings across different forms of chronic pain (23).

Since sex differences have been reported in the response to chronic pain and opioids (reviewed in (24-26). We trained male and female TetTag mice (27) to self-administer sucrose (natural reward) or oxycodone (opioid reinforcer) for 10 days followed by 7 days of forced abstinence (see timeline in Fig 1). In an initial drug seeking session on abstinence day 7, drug-seeking ensemble neurons were “tagged.” The next day we induced neuropathic pain via a spared nerve injury (SNI) and mice remained in forced abstinence until day 21 when mice were allowed a second drug-seeking session. We assessed seeking behavior, as well as electrophysiological measurements of ensemble and non-ensemble layer V/VI neurons in the PrL region of the dmPFC. This experimental design allowed us to distinguish the effects of chronic pain on opioid drug-seeking from self-administration and the associated intrinsic excitability of the ensemble neurons in the PrL. We tested the hypothesis that SNI would increase oxycodone seeking behavior even when self-administration did not occur during the pain state. In addition to this, we tested the hypothesis that an increase in drug-seeking was associated with an increase in the intrinsic excitability of the specific drug-seeking ensemble neurons.

**Figure 1.**
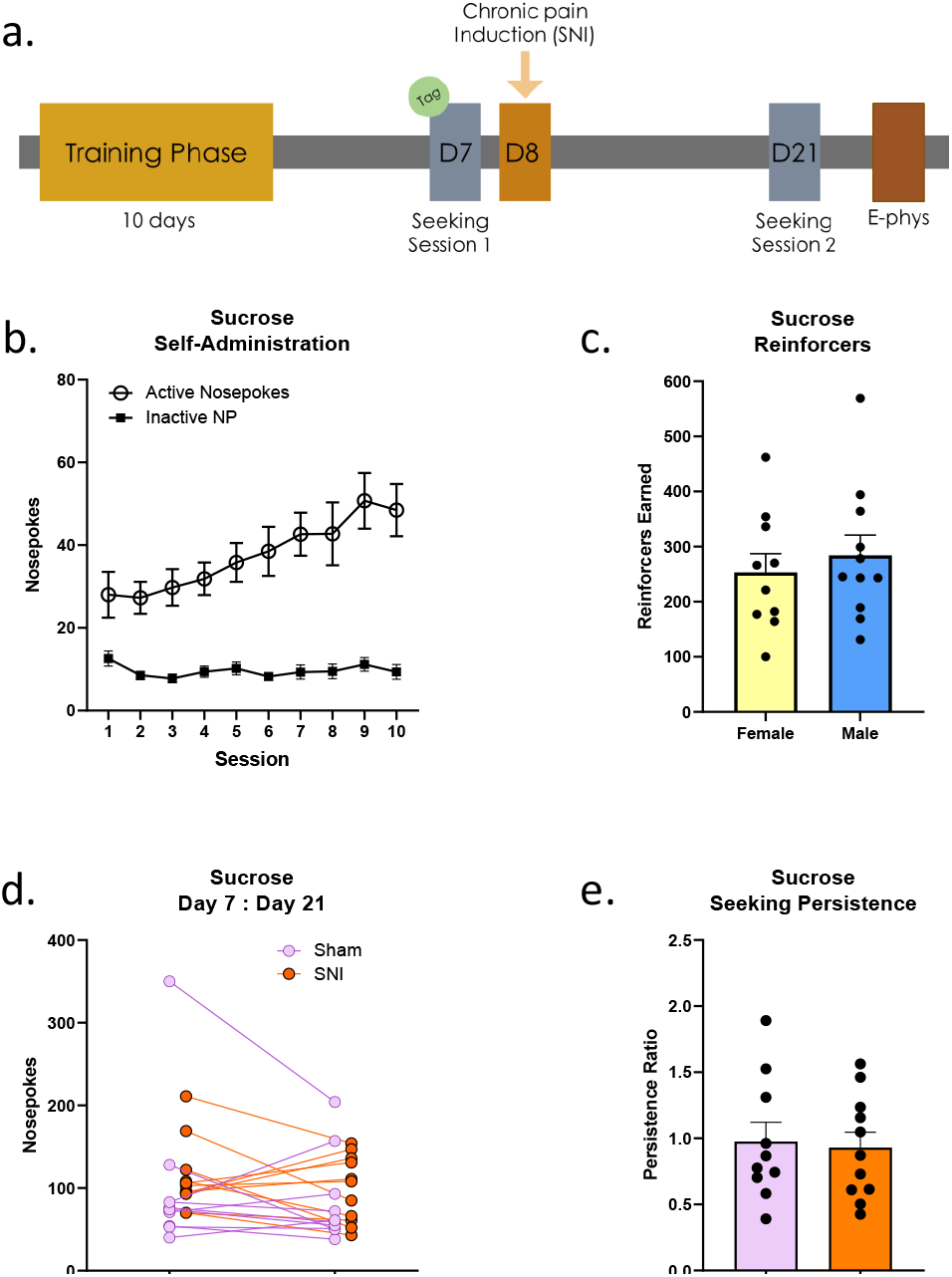
Sucrose seeking behavior following SNI is not different compared to sham. **a**. Experimental timeline completed by all mice. **b**. Self-administration sessions across 10 days. Active nosepokes denoted as open circles, inactive nosepokes denoted as black boxes. **c**. Total number of sucrose reinforcers earned by females (yellow) compared to male (blue) (t(19) = 0.609, p = 0.550). **d**. Individual values of number of active nosepokes on D7 to D21 seeking. **e**. Quantification of the persistence ratio, defined as D21/D7 active nosepokes (t(19) = 0.247, p = 0.808). Symbols/bars represent mean ± SEM.

## MATERIALS & METHODS

### Animal Model

Thirty one female and 31 male TetTag mice (c-fos-ttA/ TRE-H2B-EGFP) were used to capture specific ensemble neurons activated on day 7 seeking. Animals were bred as described in (28). When maintained on a doxycycline diet (40 ppm, Teklad Custom Research Diets, Envigo, Harlan Laboratories, Madison, WI) at libitum, very minimal tagging takes place (described in (28)). When doxycycline diet is removed and there is robust neuronal firing, the *c-fos* promoter in transgene 1 drives expression of the tetracycline transactivator (ttA) which can then bind to the tet-response element (TRE) in transgene 2 and subsequently transcribe and translate the histone H2B-EGFP fusion protein which remains stable for several weeks (27). The tagging period is closed by resuming a high dox diet (625mg/kg, Teklad Custom Research Diets, Envigo, Harlan Laboratories, Madison, WI). Mice were kept in a 12 hour reverse light cycle (0730-1930) in an AAALAC-approved facility. All animals were handled for a minimum of 3 days using procedures described in (29) prior to any surgical or behavioral experiments. All animal procedures were performed in accordance with the Medical College of Wisconsin animal care committee’s regulations and the NIH guidelines for the care and use of laboratory animals.

### Jugular Catheterization

Surgeries were performed under isoflurane (Vetequip - Akorn, Inc., Lake Forest, IL) anesthesia (3%-5% induction, 1%-3% maintenance) as described in (28,30). Tubing (micro-renathane, Braintree Scientific, Braintree, MA) was placed into the right jugular vein and secured with suture (4-0 black braided silk, non-absorbable, Henry Schein INC, Melville, NY). The cannula port was fixed beneath the hypodermis just below the interscapular region. Mice were given 5 mg/kg carprofen following surgery and triple antibiotic ointment (bacitracin zinc-neomycin sulfate-polymyxin B sulfate) at the incision sites. Post-operative measures the following day included 5 mg/kg carprofen and more triple antibiotic ointment at the incision site. On the third day following surgery and beyond, mice were given a heparin-saline (100 units/ml, Human Pharmaceuticals) flush daily to prevent clotting. One day before self-administration (SA), mice were given a Brevital (methohexital 9 mg/kg, i.v; JHP Pharmaceuticals, Rochester, NY) test to check for catheter patency before beginning the experiment. Catheter patency was confirmed by presence of a behavioral response occurring within 5 seconds of infusion.

### Experimental Apparatus (Self-Administration/Seeking Sessions)

Operant conditioning occurred in mouse-specific behavioral chambers (Med Associates, St. Albans, VT). Mice were provided with a left and right nosepoke and learned to associate one with receipt of the drug (active nosepoke side was counterbalanced). When the active nosepoke was chosen, an intravenous infusion of 0.03 mg/kg of oxycodone (Medisca, Plattsburgh, NY, USA) was dispensed (∼40 µl, ∼2.1 sec, time and volume adjusted for animal weight) while a cue light appeared simultaneously (lasting 5 seconds). Natural reward control group received 30 μl of 10% sucrose (Sigma-Aldrich, St. Louis, MO) prepared in hyperchlorinated drinking water. There was A 10 second timeout period in which no drug was administered immediately followed a drug-receiving active nosepoke, but any active nosepokes during that time were recorded. Each self-administration session lasted for 1 hour for sucrose-receiving mice and 2 or 6 hours for oxycodone-receiving mice. The session ended early if a total of 64 reinforcers were earned in the 1- and 2-hour sessions, or 192 reinforcers for the 6-hour SA sessions. In order to advance in the study, mice needed to meet three criteria: more than 10 reinforcers needed to be earned in a session, a preference of 2:1 active to inactive nosepokes must be met, and the previous listed criteria must be met for 2/3 of the last self-administration days. If the criteria were not met, mice were allowed up to 3 extra days to meet criteria. A Brevital test (methohexital 9 mg/kg, i.v.) was performed after the entirety of the SA phase to confirm catheter patency. Seeking sessions took place in the same chamber as the SA for a 2-hour duration (on day 7 and day 21 of abstinence) in which no reinforcers were earned, but the cue light was delivered in response to an active nosepoke.

### Spared Nerve Injury & Mechanical Sensitivity Assay

The spared nerve injury (SNI) was performed under isoflurane anesthesia as described for jugular catheterization and consisted of ligation and transection of the common peroneal and sural nerve branches of the sciatic nerve but sparing of the tibial nerve as similarly described (31). Vicryl suture (5-0, Ethicon, Guaynabo, Puerto Rico) was used for the muscle and VISORB polyglycolic acid suture (5/0 absorbable suture, CP Medical, Norcros, GA) was used for the skin, followed by triple antibiotic ointment at the incision site. Mice were tested for mechanical sensitivity using the von Frey assay for 2 baseline measurements prior to SNI and then post-surgery on abstinence days 9,10, 12, 15, and 20.

### Electrophysiology

Mice were first anesthetized with isoflurane and then swiftly decapitated. The brain was then mounted in a low-melting-point agarose gel (3%) to keep stable for slicing. Once chilled, it was moved to a vibratome (Leica VT1200s, Nussloch, Germany) to be sectioned in 200 µm slices at a speed of 0.16 mm/s in an NMDG-based cutting solution containing the following: 92 mM NMDG, 2.5 mM KCl, 1.25 mM NaH2PO4, 26 mM NaHCO3, 20 mM HEPES, 25 mM glucose, 2 mM thiourea, 5 mM Na-ascorbate, 3 mM Na-pyruvate, 0.5 mM CaCl2·2H2O, and 7 mM MgSO4 (pH 7.3–7.4 with HCl). Slices incubated for 20 minutes as Artificial Cerebral Spinal Fluid (ACSF, containing: 119 mM NaCl, 3 mM KCl, 2 CaCl2, 1 mM MgCl2, 1.25 mM NaH2PO4, 25 mM NaHCO3, and 10 mM glucose at room temperature) is titrated into the cutting solution in 5-minute intervals as previously described in (32). Lastly, slices were placed into ACSF to incubate for 30 minutes. All solutions were saturated with 95% O2 and 5% CO2.

Whole-cell patch-clamp recordings were made using patch-clamp amplifiers (Multiclamp 700B). Data acquisition and analysis were performed using DigiData 1550B digitizers and the analysis software pClamp 10.7 (Molecular Devices). Glass micropipettes were pulled to 4-6 Ω and filled with internal solution. Internal solution was made using 140 K-gluconate, 10 KCl, 10 HEPES, 0.2 EGTA, 2 MgCl2, 4 Mg-ATP, 0.3 Na2GTP, 10 Na2-phosphocreatine (pH 7.2 with KOH). Neurons selected (2-4 EGFP negative and 2-4 EGFP positive) were contralateral to the injury side in layer V/VI of the PrL (Bregma 2.7-3.7). Each neuron was recorded in current clamp mode, which included measurement of action potentials (AP) in response to current injections (−60 to 120 pA, 20 pA steps), rheobase at 5 pA steps, and input resistance (IR) measured in -20 pA steps. Resting membrane potential was also collected.

### Statistics

T-tests were used to compare total number of reinforcers earned between males and females after SA, and again when comparing the persistence ratio of sham and SNI animals. Generalized linear mixed effect models were used to analyze the association of injury and cell type (EGFP-/+) with measures of intrinsic excitability (AP AUC, rheobase, IR). The mixed effect model, in addition to measuring fixed effects, also adjusts for correlations for measures within animal (and within neuron input-output AP firing).

Compound symmetry structures were used for the mixed effects models. Modeling assumptions were rigorously checked, including behavior of residuals, to ensure no major departures from homoscedasticity or normality assumptions. Multiplicity adjustments, wherever relevant, were performed using Tukey’s procedure.

## RESULTS

### Sucrose-seeking in mice is not influenced by SNI

In order to answer our larger question of how neuropathic pain may affect reactivity to drug-related stimuli, we first started our experiments looking at sucrose as a natural reward. Across the 10 days of 1-hour sucrose self-administration (Fig 1a and 1b), the total number of reinforcers earned between females and males was not significantly different (Fig 1c; t(19) = 0.609, p = 0.550) therefore subsequent outcome measures were pooled between sexes. Mice were counterbalanced to either sham or SNI based on their D7 seeking. When we compared the number of nosepokes from D7 seeking (before SNI) to D21 seeking (after SNI) as measured by a seeking persistence ratio 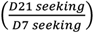, we discovered no significant difference among mice that received SNI compared to sham (Fig 1e; t(19) = 0.247, p = 0.808).

### Specific sucrose-seeking ensemble neurons show more intrinsic excitability compared to non-ensemble neurons

We hypothesized that sucrose-seeking ensemble neurons would exhibit higher intrinsic excitability compared to non-ensemble neurons, as acute sucrose seeking ensemble hyperexcitability has been identified in the dmPFC of *c-fos*-EGFP mice (33). We measured input-output curves by recording action potentials (AP) evoked by 20 pA step current injections in both ensemble and non-ensemble neurons of sham and SNI animals (Fig 2a, 2a’). When plotted as an area-under the curve (AUC), there was a main effect of ensemble neurons exhibiting higher intrinsic excitability when compared to non-ensemble neurons (Fig 2b, 2b’; p = 0.003) but no main or interaction effect of SNI (p = 0.965, p = 0.208). We also noted a main effect in the intrinsic excitability of ensemble neurons versus non-ensemble neurons with respect to rheobase and input resistance (Fig 2c, 2c’ and 2d, 2d’; p = 0.005 and p = 0.021,). Neurons from SNI mice did not show a significant difference in rheobase and input resistance when compared to sham (Fig 2c, 2c’ and 2d, 2dc’; p = 0.744 and p = 0.955). There were also no significant differences in resting membrane potential across injury or celltype (p = 0.257 and p = 0.090). Taken together, there was no effect of SNI on sucrose seeking or intrinsic excitability of PrL pyramidal neurons, but sucrose seeking ensemble neurons exhibited higher intrinsic excitability when compared to non-ensemble neurons in mice that received SNI or sham injury.

**Figure 2.**
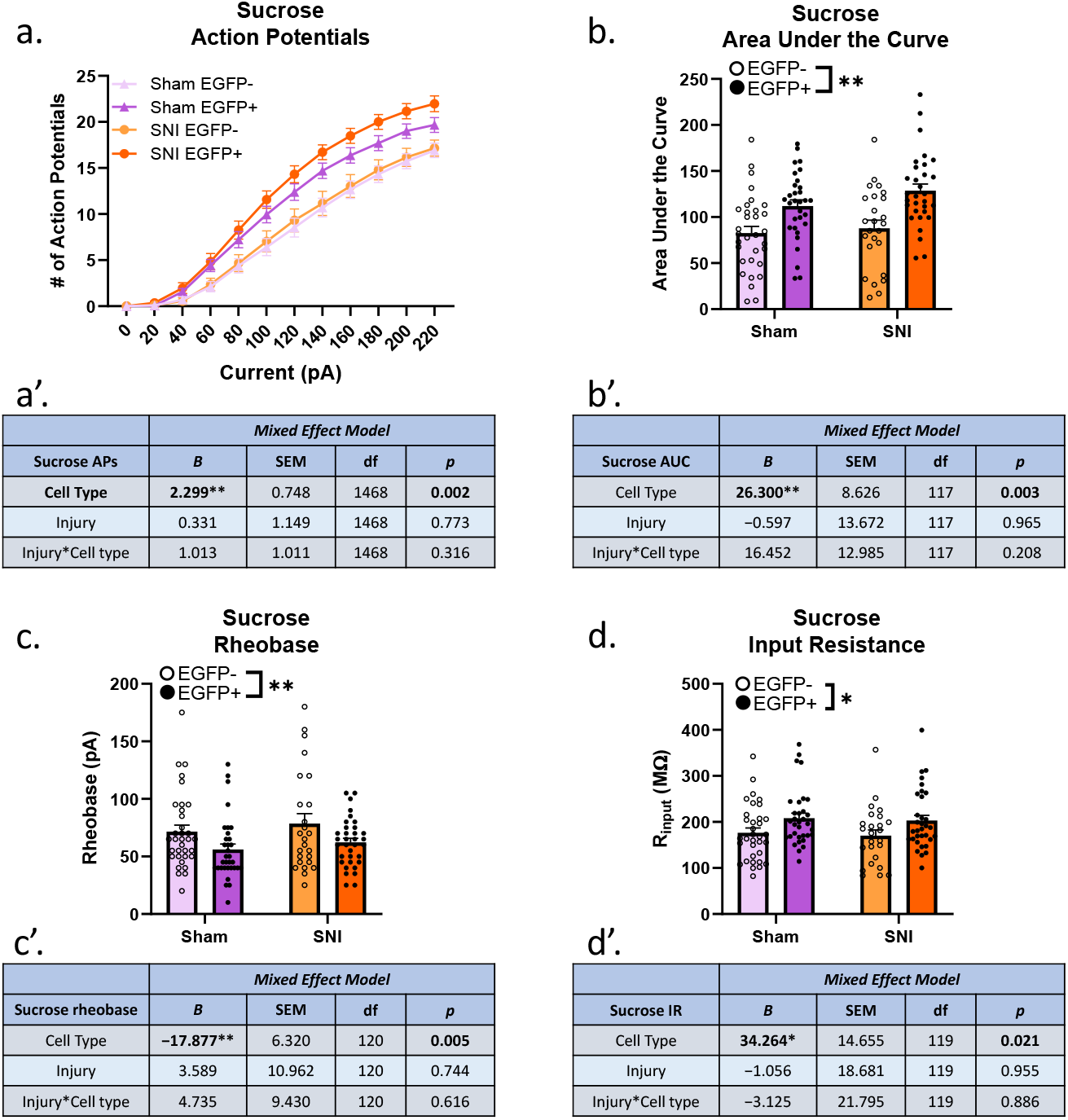
Sucrose-seeking PrL layer V/VI ensemble neurons are significantly more excitable compared to non-ensemble neurons. **a**. Input-output curve showing action potentials recorded given 20 pA current injection steps in both sham and SNI mice and their respective EGFP-vs EGFP+ neurons. All statistical analyses accounted for multiple neurons coming from an animal. **b**. Plotted area under the curve, per neuron, illustrating differences between EGFP- and EGFP+ neurons within sham and SNI. Statistical analyses accounted for multiple neurons coming from an animal. **c**. Current injections increased in 5 pA increments until the first action potential was recorded for the rheobase. **d**. Input resistance was measured in decreasing -20 pA steps. Symbols/bars represent mean ± SEM. *p<0.05, **p<0.01.

### 2-hour oxycodone self-administration sessions did not yield behavioral differences in seeking upon induction of SNI

Mice self-administered oxycodone across the 10 days during 2-hour sessions, and there was no significant difference in the total number of reinforcers earned between males and females (Fig 3a and 3b; t(11) = 1.23, p = 0.244). Therefore, males and females were pooled for subsequent outcome measures. We then compared D7 seeking to D21 and showed there was no significant difference in mice that received SNI compared to sham (Fig 3c and 3d; t(11) = 0.784, p = 0.686).

**Figure 3.**
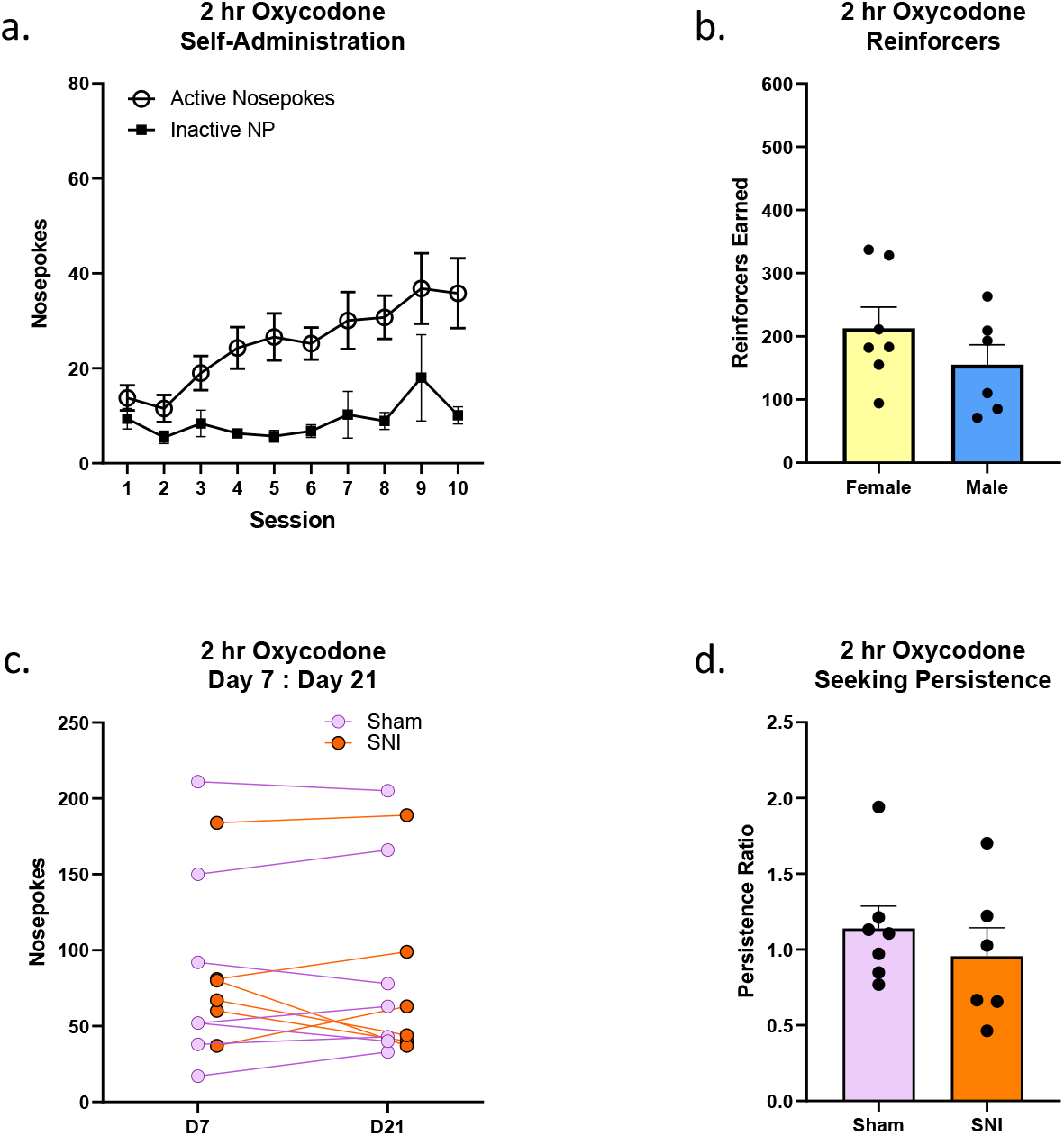
2hr oxycodone seeking behavior following SNI is not different compared to sham. **a**. Self-administration sessions across 10 days. Active nosepokes denoted as open circles, inactive nosepokes denoted as black boxes. **b**. Total number of sucrose reinforcers earned by females (yellow) compared to male (blue) (t(11) = 1.231, p = 0.244). **c**. Individual values of number of active nosepokes on D7 to D21 seeking. **d**. Quantification of the persistence ratio, defined as D21/D7 active nosepokes (t(11) = 0.784, p = 0.4496). Symbols/bars represent mean ± SEM.

### SNI did not affect intrinsic excitability of ensemble neurons in 2-hour oxycodone SA mice

We further investigated the electrophysiology of PrL pyramidal neurons given 20 pA current injections and discovered an interaction effect of injury and ensembles on action potentials (Fig 4a, 4a’; p = 0.004), and a main effect of ensemble membership (Fig 4a, 4a’; p = 0.048). However, we found no main effect of injury with regards to intrinsic excitability when quantified as area under the curve (Fig 4b, 4b’; p = 0.358), and no significant difference in excitability of ensemble neurons or interaction with injury (Fig 4b, 4b’; p = 0.640, p = 0.365). There was also no effect of SNI or ensemble membership in measures of rheobase (Fig 4c, 4c’; p = 0.662, p = 0.316) or input resistance (Fig 4d, 4d’; p = 0.996 and p = 0.956). There was no difference in resting membrane potential across injury or celltype (p = 0.780 and p = 0.580). Taken together, the mice in the 2-hour oxycodone self-administration group did not exhibit SNI-associated changes in drug seeking, and only showed an SNI-associated increase in intrinsic excitability with respect to area under the curve from the input-output recordings of action potentials.

**Figure 4.**
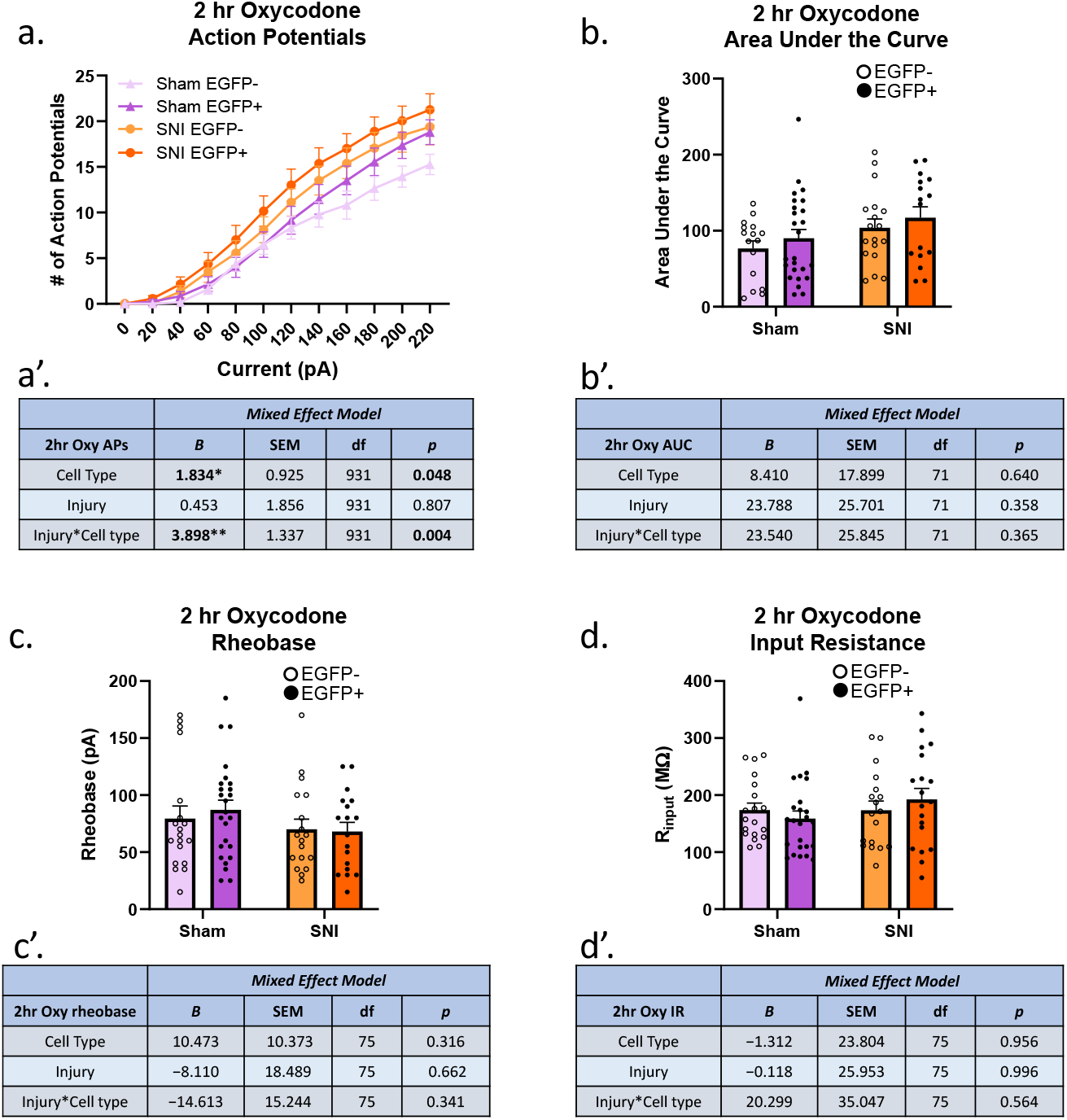
2hr oxycodone-seeking PrL layer V/VI ensemble neurons in mice that received SNI fire more action potentials in response to current injections compared to sham. **a**. Input-output curve showing action potentials recorded given 20 pA current injection steps in both sham and SNI mice and their respective EGFP-vs EGFP+ neurons. All statistical analyses accounted for multiple neurons coming from an animal. **b**. Plotted area under the curve, per neuron, illustrating differences between EGFP- and EGFP+ neurons within sham and SNI. **c**. Current injections increased in 5 pA increments until the first action potential was recorded for the rheobase. **d**. Input resistance was measured in decreasing -20 pA steps.

### There were sex differences in total consumption and subsequent seeking behavior in response to SNI in mice that had 6-hour oxycodone self-administration sessions

Extended access drug self-administration models are proposed to better model compulsive drug use in humans (34,35). Using the same timeline as previously described (Fig 1a), male and female mice learned to self-administer oxycodone across the 10-day training period (Fig 5a), but this time for 6-hour sessions. Across all SA sessions, the total number of reinforcers earned per animal was calculated. There was a significant difference in the total summation of reinforcers following self-administration, such that females earned nearly 2-fold reinforcers of oxycodone than males (Fig 5b; t(26) = 4.07, p = 0.0004). Due to this difference, subsequent outcome measures were analyzed separately for each sex. D7 seeking values were counterbalanced for assignment to sham or SNI, and individual measurements in seeking behavior were plotted (Fig 5c and 5d). When quantified as a persistence ratio, interestingly, female mice that were given SNI following D7 seeking significantly increased their seeking behavior on D21 (Fig 5e; t(10) = 3.553, p = 0.0052). When we examined the male seeking behavior, we saw no significant difference in male mice assigned to the SNI group (Fig 5f; t(12) = 0.421, p = 0.681). There was no significant association between intake and persistence ratio (r^2^ = 0.002, p = 0.892), suggesting that the sex difference in intake did not contribute to the SNI-increased oxycodone seeking. Thus, chronic pain induced by SNI increased seeking 2 weeks later, specifically in females but not male mice.

**Figure 5.**
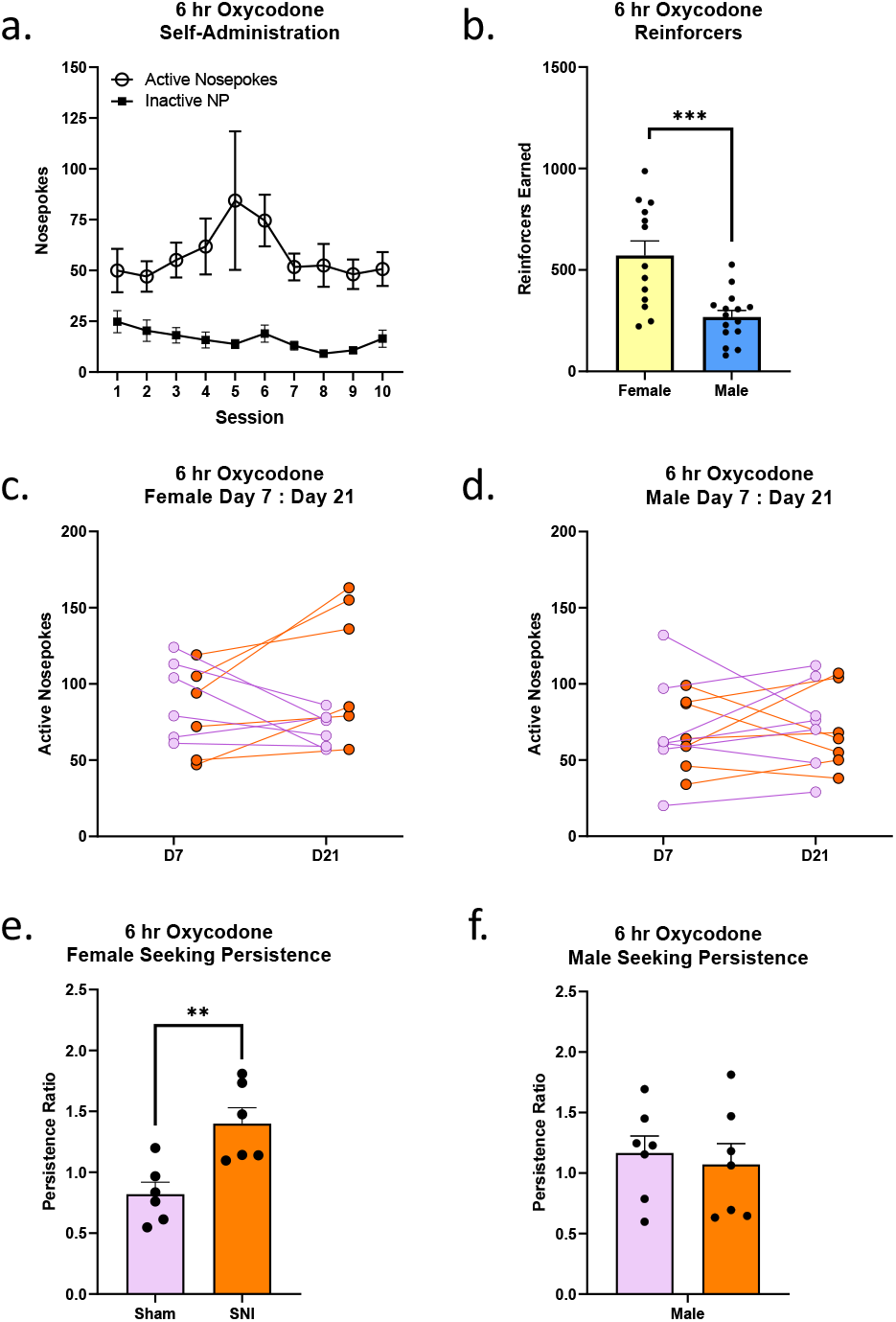
6hr oxycodone self-administration in females is significantly different compared to males, and females show a significant increase in seeking behavior following SNI. **a**. Self-administration sessions across 10 days. Active nosepokes denoted as open circles, inactive nosepokes denoted as black boxes. **b**. Total number of sucrose reinforcers earned by females (yellow) compared to male (blue) (t(26) = 4.07, p = 0.0004). **c**. Individual values of number of active nosepokes on D7 to D21 seeking in the female mice. **d**. Individual values of number of active nosepokes on D7 to D21 seeking in the male mice. **e**. Female seeking persistence ratio, defined as D21/D7 active nosepokes (t(10) = 3.553, p = 0.0052. **f**. Male seeking persistence ratio (t(12) = 0.421, p = 0.681). Symbols/bars represent mean ± SEM. **p<0.01, ***p<0.001.

### 6-hour oxycodone SA females exhibited a significant increase in PrL intrinsic excitability in response to SNI

We then investigated our hypothesis at the neuronal level, postulating that an increase in intrinsic excitability specifically within the ensemble neurons could be responsible, in part, for an increase in behavioral seeking behavior in females. In female mice, we found that there was a significant main effect of SNI on the intrinsic excitability, such that the SNI neurons were had greater action potential firing in response to current injections when compared to sham (Fig 6a, 6a’; p < 0.001). However, the significance seen was largely driven by the non-ensemble neurons and is not *specific* to an increase in ensemble intrinsic excitability (see AUC, Fig 6b, 6b’). An interaction effect of injury*celltype was also discovered in the number of APs fired (Fig 6a, 6a’; p < 0.001). Additionally, significant differences were also detected as a main effect among ensemble versus non-ensemble neurons with respect to rheobase (Fig 6c, 6c’; p < 0.001) and input resistance (Fig 6d, 6d’; p < 0.001). There was no significant difference in the resting membrane potential of neurons recorded from across injury or ensemble membership (p = 0.598 and p = 0.363). Given the increase in intrinsic excitability seen with respect to the APs, we wanted to see if there was an association with an increase in seeking as measured by the persistence ratio, so we plotted the persistence ratio against the AUC value using a linear mixed effect regression. While there was no significant association from injury, we *did* detect a significant difference of persist_ratio*celltype (Fig 6e, 6e’; p = 0.001), such that an increase in intrinsic excitability of cell type showed a relationship with an increase in seeking persistence. There was no main effect of injury.

**Figure 6.**
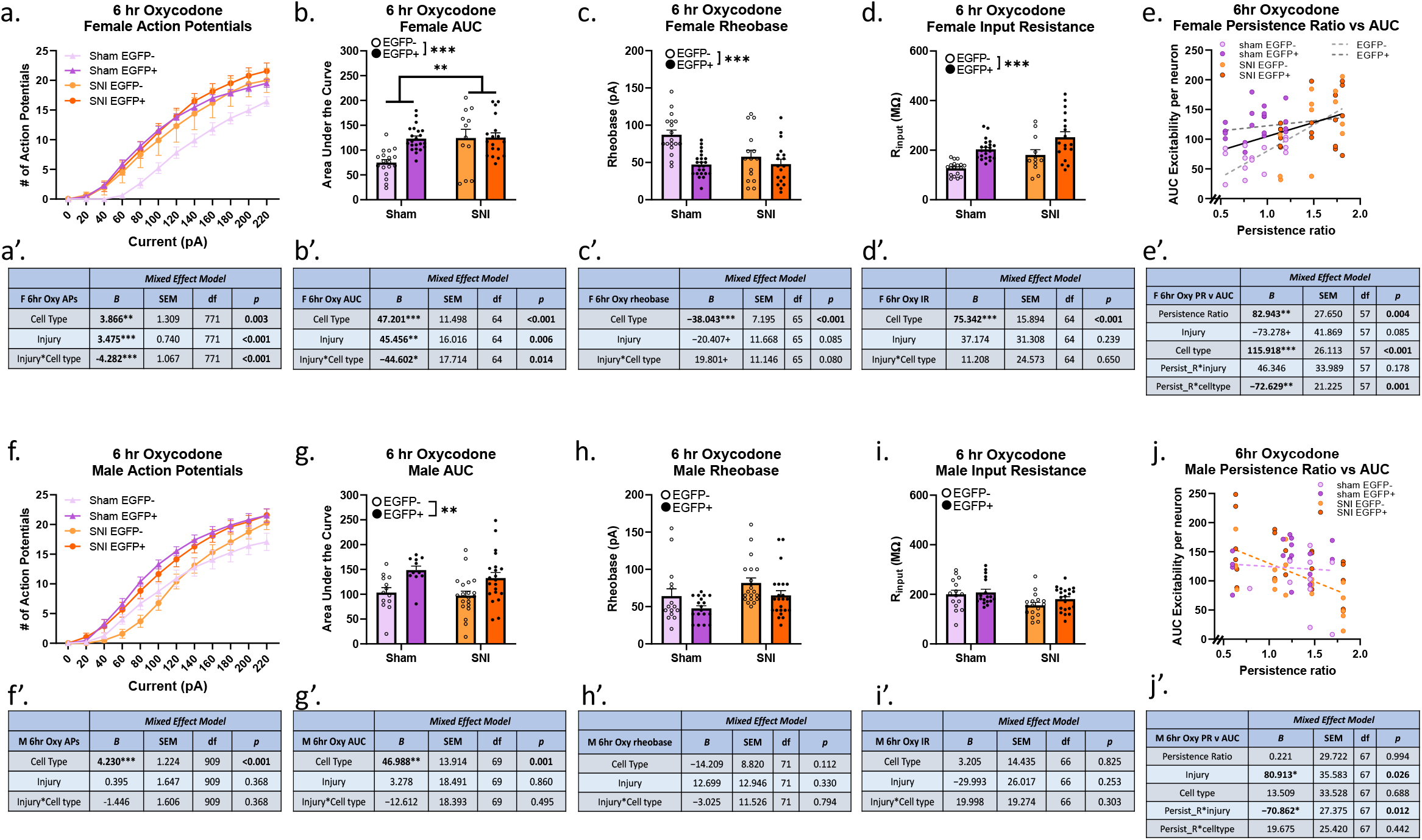
6hr Oxycodone-seeking PrL layer V/VI neurons in female mice that received SNI showed hyperexcitability as a result of SNI induction and previous SA history, but not in male mice. **a & f**. Input-output curve showing action potentials recorded given 20 pA current injection steps in both sham and SNI mice and their respective EGFP-vs EGFP+ neurons. All statistical analyses accounted for multiple neurons coming from an animal. **b & g**. Plotted area under the curve, per neuron, illustrating differences between EGFP- and EGFP+ neurons within sham and SNI. **c & h**. Current injections increased in 5 pA increments until the first action potential was recorded for the rheobase. **d & i**. Input resistance was measured in decreasing -20 pA steps. **e & j**. AUC per neuron plotted against persistence ratio. Symbols/bars represent mean ± SEM. *p<0.05 **p<0.01, ***p<0.001.

In male mice, when we recorded the number of action potentials following 20 pA steps and plotted AUC, we noted no significant difference of SNI mice when compared to sham in the number of action potentials (Fig 6f, 6f’; (APs) p = 0.368 and Fig 6g, 6g’; (AUC) p = 0.860), however we did detect a significant difference in the ensemble neuron cell excitability compared to non-ensemble neurons (Fig 6f, 6f’; (APs) p < 0.001 and Fig 6g, 6g’; (AUC) p = 0.001). There was no significant change in cell excitability comparing SNI to sham with respect to rheobase and input resistance (Fig 6h, 6h’; p = 0.330 and Fig 6i, 6i’; p = 0.253), nor for ensemble membership (Fig 6h, 6h’; p = 0.112 and Fig 6i, 6i’; p = 0.825). Resting membrane potential yielded no significant differences across injury or celltype (p = 0.363 and p = 0.570). Despite a lack of significance in behavioral changes or electrophysiological changes resulting from SNI, we still examined the relationship between persistence ratio and AUC value as we did in the females and discovered that there was in fact a significant association across injury, such that mice with a higher persistence ratio yielded a significant relationship with a decrease in neuron AP AUC values (Fig 6j, 6j’; p = 0.012). This association was not observed in relation to cell type.

Taken together, when female mice received SNI after their initial drug seeking session, two weeks later we saw an increase in seeking behavior; this was not seen in the male mice. When we recorded from PrL pyramidal neurons in females, we discovered that neurons from SNI mice exhibited higher intrinsic excitability (as measured by APs and AP AUC,) than sham, and this difference was largely driven by the non-ensemble neurons. When we recorded from PrL pyramidal neurons in males, we saw a significant difference in ensemble neurons compared to non-ensemble neurons with respect to APs and AP AUC, but there was no significant difference in neurons recorded from SNI mice compared to sham across any of the measures of intrinsic excitability.

## DISCUSSION

Here, we examined the effects of neuropathic pain on natural and drug reward seeking behavior and neuronal excitability in the PrL. Using the TetTag mouse model we captured D7 seeking ensemble neurons and compared intrinsic excitability of these neurons to non-ensemble neurons. We demonstrated that mice that self-administered sucrose or short-access (2 hour) oxycodone did not exhibit behavioral changes in seeking when SNI was induced after the initial seeking session. In mice that had long-access (6 hour) oxycodone self-administration, SNI induced during abstinence elevated oxycodone seeking in females, but not males. In 6-hour oxycodone females, intrinsic excitability (as measured by current-spike relationship) of non-ensemble PrL pyramidal neurons in females was elevated by SNI, and excitability of PrL pyramidal neurons correlated with oxycodone seeking in female mice. The increased excitability of PrL non-ensemble cells, positive association between excitability and oxycodone seeking, and lack of either behavioral or electrophysiological changes by SNI in males suggests that SNI may have expanded the size of the dmPFC oxycodone seeking ensemble specifically in females. Pain-associated ensembles that are necessary for driving pain-associated changes in affect have been identified in the basolateral amygdala and dmPFC (36,37). The dmPFC pain ensemble was found to project to regions important in opioid seeking, including the nucleus accumbens and central amygdala (36), suggesting that this ensemble could overlap with a previously established drug seeking ensemble. Additionally, longitudinal calcium imaging studies within the PrL region of the dmPFC have shown that weakly tuned neurons at early behavioral time points can be recruited into ensembles that they showed an early bias toward (38).

Many studies have examined the comorbidity of pain and drug-seeking behavior through the lens of opioids as a negative reinforcer in which pain precedes the administration of the analgesic (21,22,39,40). In the context of acute pain, Shaham and colleagues found that intraplantar capsaicin did not cause reinstatement of fentanyl seeking, nor did intraperitoneal injections of lactic acid for heroin seeking (41). However, Hipólito and colleagues showed that in an inflammatory pain state from CFA (Complete Freund’s Adjuvant) intraplanar injections, high doses of heroin resulted in significantly more drug self-administration and the motivated behavior was maintained (as measured by progressive ratio), but this was not seen in the low dose heroin group (21). Additionally, CFA has been shown to have long-lasting affective effects, and after a single CFA injection followed by 11 days of morphine self-administration (0.2 mg/kg/infusion), morphine intake was not affected (22). However, after 13 days of abstinence, rats showed elevated drug seeking and did not extinguish lever-pressing beyond 5 days. Similarly, SNI did not affect self-administration of 0.1 mg/kg/infusion morphine, but led to persistent increases in drug seeking during extinction testing (42). Our current study was designed to isolate the pain state from opioid self-administration. With the pain-associated negative reinforcement of opioid self-administration removed, we interpret our findings to indicate that in females, chronic pain-associated increases in opioid seeking can occur in the absence of opioid alleviation of pain.

Neuropathic pain leads to neuronal and synaptic plasticity within the brain and has both short- and long-term effects at the circuit level. Alterations in limbic circuit plasticity have been observed up to 12 months post-SNI and exhibit in mice (43). While it is widely understood that the PFC projects to the nucleus accumbens (NAc) in reward circuitry, this projection has also been shown to play a role during a shift from acute to chronic pain (44). Work by Guida and colleagues showed that SNI mice measured at 1 and 12 months out from injury exhibited impaired long-term-potentiation (LTP) at this pathway, which they hypothesize may play a role in the affective experience of pain (43). Furthermore, dopaminergic projections from the ventral tegmental area (VTA) to the mPFC have been shown to play a role in the chronic pain, and optogenetic stimulation of this projection alleviated mechanical allodynia and produced a conditioned place preference (45). VTA dopaminergic neurons in a pain state contributes to anhedonia-like behavior by way of decreased intrinsic excitability of VTA DA neurons and a reduction in mesolimbic dopamine release (46). At the cellular level, prior work in rats 2 weeks out from SNI reported layer II/III pyramidal neurons in the PrL had an increase in intrinsic excitability (20). In mice, electrophysiological recordings from layer V pyramidal neurons within the PrL following SNI showed an acute peak of hyperexcitability within the first week, and then became significantly hypoexcitable at 35 days out from SNI (47). It is unknown if prior opioid exposure changes this phenomenon. Considering our observed differences in intrinsic excitability in PrL pyramidal neurons recorded from mice with sucrose or 6-hour oxycodone self-administration, opioid exposure may delay the transition from hyper-to hypo-excitability of these cells.

Memories rely on ensemble neurons, which can be strengthened in particular brain regions when paired with a substance reinforcer (48,49). The specificity of ensembles has been described to play a role in context-induced relapse in rats that previously self-administered heroin (49), as well as work done in cocaine (48,50). The strength and population of an ensemble can grow following substance self-administration, and this has been shown through behavioral work and electrophysiological studies (50-52). Therefore, we hypothesized that the addition of neuropathic pain could perhaps further enhance the neuronal ensemble intrinsic excitability within the PFC.

Interestingly, we saw increases in intrinsic excitability in female mice that received SNI, but this effect was isolated to non-ensemble neurons. Further analysis indicated that there was a positive association between seeking persistence and AP AUC in female, but not male mice. In males, we did not see a significant change in intrinsic excitability of sham versus SNI neurons with respect to rheobase or input resistance. We did, however, discover a relationship of higher persistence ratios in males being associated with a decrease in intrinsic excitability. While this is not what we initially hypothesized to observe, SNI-associated hypoexcitability has been shown to occur in PFC pyramidal neurons at more chronic durations than tested here (37). In our paradigm, we found that this effect was specific to males that self-administered oxycodone compared to sucrose. Thus, we identified contingent sex differences (26) where oxycodone SA (and abstinence) interacted with SNI in a sex-dependent manner in both behavioral and electrophysiological measures, even though oxycodone and SNI were never present at the same time. While we were able to correlate our behavioral outcomes with the electrophysiological recordings in 6-hour oxycodone females, we can not elucidate the mechanism driving seeking in the females given our current design. Future studies are needed to determine if SNI-associated changes in oxycodone seeking and PrL excitability evolve over time, and whether the SNI-associated PrL excitability is causally linked to elevated oxycodone seeking.

The present studies have some limitations due to experimental design. The motivational component was important to our experiment, but we were not able to control for the total number of reinforcers earned during self-administration. It is evident that males in the 6-hour sessions did not self-administer the same number of reinforcers as the females did, and this difference in total amount of oxycodone taken could influence future seeking behavior or the physiological response to SNI. Correlations examining the relationship between intake and persistence ratio in this study did not yield a significant relationship, suggesting that intake differences do not explain sex differences in seeking. Another potential limitation of this experiment is that there may be a ceiling effect in neuron excitability, and the selective increase in non-ensemble neurons in female SNI mice may be an artifact of ensemble neurons already having a maximal current-spike response. Furthermore, we used the von Frey assay to measure mechanical withdrawal throughout our study to validate maintained hyperalgesia. This measure is reflexive in nature and does not address the affective dimension of pain (53), which is largely regulated by the PFC (54,55). Therefore, assessment of pain behavior is limited to the sensory component. Finally, our measurements were only performed at a single timepoint. Future studies are needed to determine if the SNI-associated increase in oxycodone seeking (and associated neuroplasticity) occurs at more chronic timepoints.

## Conclusions

In conclusion, we find that chronic pain can increase oxycodone seeking in females even when drug intake preceded the pain state. Pain is reported to be a trigger for drug craving and relapse in humans (56,57), and prior substance misuse is associated with greater opioid misuse in patients with chronic pain. History of treatment in a drug or alcohol rehabilitation facility had a 93% predictive validity in determining which patients had substance misuse (58), while low dose opioid therapy did not elevate risk of developing opioid misuse in chronic pain patients without prior substance use disorder (59). Thus, our data from females are consistent with clinical reports that chronic pain can promote drug craving and relapse and support the hypothesis that chronic pain may lead to neuroadaptations which promote drug seeking.

## Abbreviations

SA: self-administration
SNI: spared nerve injury

## REFERENCES

1. Centers for Disease Control and Prevention, Opioid Overdose.

2. Mason, M., Welch, S. B., Arunkumar, P., Post, L. A., and Feinglass, J. M. (2021) Notes from the Field: Opioid Overdose Deaths Before, During, and After an 11-Week COVID-19 Stay-at-Home Order - Cook County, Illinois, January 1, 2018-October 6, 2020. MMWR Morb Mortal Wkly Rep 70, 362–363

3. Centers for Disease Control and Prevention (2020), Increase in Fatal Drug Overdoses Across the United States Driven by Synthetic Opioids Before and During the COVID-19 Pandemic., December 17, 2020 Ed.

4. Edlund, M. J., Martin, B. C., Russo, J. E., DeVries, A., Braden, J. B., and Sullivan, M. D. (2014) The role of opioid prescription in incident opioid abuse and dependence among individuals with chronic noncancer pain: the role of opioid prescription. Clin J Pain 30, 557–564

5. Dahlhamer, J., Lucas, J., Zelaya, C., Nahin, R., Mackey, S., DeBar, L., Kerns, R., Von Korff, M., Porter, L., and Helmick, C. (2018) Prevalence of Chronic Pain and High-Impact Chronic Pain Among Adults - United States, 2016. MMWR Morb Mortal Wkly Rep 67, 1001–1006

6. Volkow, N. D., and McLellan, A. T. (2016) Opioid Abuse in Chronic Pain--Misconceptions and Mitigation Strategies. N Engl J Med 374, 1253–1263

7. Kyranou, M., and Puntillo, K. (2012) The transition from acute to chronic pain: might intensive care unit patients be at risk? Ann Intensive Care 2, 36

8. Vowles, K. E., McEntee, M. L., Julnes, P. S., Frohe, T., Ney, J. P., and van der Goes, D. N. (2015) Rates of opioid misuse, abuse, and addiction in chronic pain: a systematic review and data synthesis. Pain 156, 569–576

9. Sinha, R. (2008) Chronic stress, drug use, and vulnerability to addiction. Ann N Y Acad Sci 1141, 105–130

10. LeCocq, M. R., Randall, P. A., Besheer, J., and Chaudhri, N. (2020) Considering Drug-Associated Contexts in Substance Use Disorders and Treatment Development. Neurotherapeutics 17, 43–54

11. Stewart, J. (2008) Review. Psychological and neural mechanisms of relapse. Philos Trans R Soc Lond B Biol Sci 363, 3147–3158

12. Crombag, H. S., Bossert, J. M., Koya, E., and Shaham, Y. (2008) Review. Context-induced relapse to drug seeking: a review. Philos Trans R Soc Lond B Biol Sci 363, 3233–3243

13. Volkow, N. D., and Morales, M. (2015) The Brain on Drugs: From Reward to Addiction. Cell 162, 712–725

14. Johansen, J. P., Fields, H. L., and Manning, B. H. (2001) The affective component of pain in rodents: direct evidence for a contribution of the anterior cingulate cortex. Proc Natl Acad Sci U S A 98, 8077–8082

15. Baliki, M. N., Chialvo, D. R., Geha, P. Y., Levy, R. M., Harden, R. N., Parrish, T. B., and Apkarian, A. V. (2006) Chronic pain and the emotional brain: specific brain activity associated with spontaneous fluctuations of intensity of chronic back pain. J Neurosci 26, 12165–12173

16. Sortman, B. W., Gobin, C., Rakela, S., Cerci, B., and Warren, B. L. (2022) Prelimbic Ensembles Mediate Cocaine Seeking After Behavioral Acquisition and Once Rats Are Well-Trained. Front Behav Neurosci 16, 920667

17. Warren, B. L., Kane, L., Venniro, M., Selvam, P., Quintana-Feliciano, R., Mendoza, M. P., Madangopal, R., Komer, L., Whitaker, L. R., Rubio, F. J., Bossert, J. M., Caprioli, D., Shaham, Y., and Hope, B. T. (2019) Separate vmPFC Ensembles Control Cocaine Self-Administration Versus Extinction in Rats. J Neurosci 39, 7394–7407

18. Warren, B. L., Mendoza, M. P., Cruz, F. C., Leao, R. M., Caprioli, D., Rubio, F. J., Whitaker, L. R., McPherson, K. B., Bossert, J. M., Shaham, Y., and Hope, B. T. (2016) Distinct Fos-Expressing Neuronal Ensembles in the Ventromedial Prefrontal Cortex Mediate Food Reward and Extinction Memories. J Neurosci 36, 6691–6703

19. Tonegawa, S., Morrissey, M. D., and Kitamura, T. (2018) The role of engram cells in the systems consolidation of memory. Nat Rev Neurosci 19, 485–498

20. Cordeiro Matos, S., Zhang, Z., and Seguela, P. (2015) Peripheral Neuropathy Induces HCN Channel Dysfunction in Pyramidal Neurons of the Medial Prefrontal Cortex. J Neurosci 35, 13244–13256

21. Hipolito, L., Wilson-Poe, A., Campos-Jurado, Y., Zhong, E., Gonzalez-Romero, J., Virag, L., Whittington, R., Comer, S. D., Carlton, S. M., Walker, B. M., Bruchas, M. R., and Moron, J. A. (2015) Inflammatory Pain Promotes Increased Opioid Self-Administration: Role of Dysregulated Ventral Tegmental Area mu Opioid Receptors. J Neurosci 35, 12217–12231

22. Hou, Y. Y., Cai, Y. Q., and Pan, Z. Z. (2015) Persistent pain maintains morphine-seeking behavior after morphine withdrawal through reduced MeCP2 repression of GluA1 in rat central amygdala. J Neurosci 35, 3689–3700

23. Casale, R., Atzeni, F., Bazzichi, L., Beretta, G., Costantini, E., Sacerdote, P., and Tassorelli, C. (2021) Pain in Women: A Perspective Review on a Relevant Clinical Issue that Deserves Prioritization. Pain Ther 10, 287–314

24. Shansky, R. M., and Murphy, A. Z. (2021) Considering sex as a biological variable will require a global shift in science culture. Nat Neurosci 24, 457–464

25. Mogil, J. S. (2020) Qualitative sex differences in pain processing: emerging evidence of a biased literature. Nat Rev Neurosci 21, 353–365

26. Becker, J. B., and Chartoff, E. (2019) Sex differences in neural mechanisms mediating reward and addiction. Neuropsychopharmacology 44, 166–183

27. Tayler, K. K., Tanaka, K. Z., Reijmers, L. G., and Wiltgen, B. J. (2013) Reactivation of neural ensembles during the retrieval of recent and remote memory. Curr Biol 23, 99–106

28. Jessen, K., Slaker Bennett, M. L., Liu, S., and Olsen, C. M. (2022) Comparison of prefrontal cortex sucrose seeking ensembles engaged in multiple seeking sessions: Context is key. J Neurosci Res 100, 1008–1029

29. Olsen, C. M., and Winder, D. G. (2010) Operant sensation seeking in the mouse. J Vis Exp

30. Muelbl, M. J., Nawarawong, N. N., Clancy, P. T., Nettesheim, C. E., Lim, Y. W., and Olsen, C. M. (2016) Responses to drugs of abuse and non-drug rewards in leptin deficient ob/ob mice. Psychopharmacology (Berl) 233, 2799–2811

31. Decosterd, I., and Woolf, C. J. (2000) Spared nerve injury: an animal model of persistent peripheral neuropathic pain. Pain 87, 149–158

32. Ting, J. T., Lee, B. R., Chong, P., Soler-Llavina, G., Cobbs, C., Koch, C., Zeng, H., and Lein, E. (2018) Preparation of Acute Brain Slices Using an Optimized N-Methyl-D-glucamine Protective Recovery Method. J Vis Exp

33. Brebner, L. S., Ziminski, J. J., Margetts-Smith, G., Sieburg, M. C., Reeve, H. M., Nowotny, T., Hirrlinger, J., Heintz, T. G., Lagnado, L., Kato, S., Kobayashi, K., Ramsey, L. A., Hall, C. N., Crombag, H. S., and Koya, E. (2020) The Emergence of a Stable Neuronal Ensemble from a Wider Pool of Activated Neurons in the Dorsal Medial Prefrontal Cortex during Appetitive Learning in Mice. J Neurosci 40, 395–410

34. Koob, G. F. (2020) Neurobiology of Opioid Addiction: Opponent Process, Hyperkatifeia, and Negative Reinforcement. Biol Psychiatry 87, 44–53

35. Koob, G. F. (2009) Neurobiological substrates for the dark side of compulsivity in addiction. Neuropharmacology 56 Suppl 1, 18–31

36. Qi, X., Cui, K., Zhang, Y., Wang, L., Tong, J., Sun, W., Shao, S., Wang, J., Wang, C., Sun, X., Xiao, L., Xi, K., Cui, S., Liu, F., Ma, L., Zheng, J., Yi, M., and Wan, Y. (2022) A nociceptive neuronal ensemble in the dorsomedial prefrontal cortex underlies pain chronicity. Cell Rep 41, 111833

37. Corder, G., Ahanonu, B., Grewe, B. F., Wang, D., Schnitzer, M. J., and Scherrer, G. (2019) An amygdalar neural ensemble that encodes the unpleasantness of pain. Science 363, 276–281

38. Zhang, Y., Denman, A. J., Liang, B., Werner, C. T., Beacher, N. J., Chen, R., Li, Y., Shaham, Y., Barbera, G., and Lin, D. T. (2022) Detailed mapping of behavior reveals the formation of prelimbic neural ensembles across operant learning. Neuron 110, 674–685 e676

39. Wade, C. L., Krumenacher, P., Kitto, K. F., Peterson, C. D., Wilcox, G. L., and Fairbanks, C. A. (2013) Effect of chronic pain on fentanyl self-administration in mice. PLoS One 8, e79239

40. Martin, T. J., Kim, S. A., Buechler, N. L., Porreca, F., and Eisenach, J. C. (2007) Opioid self-administration in the nerve-injured rat: relevance of antiallodynic effects to drug consumption and effects of intrathecal analgesics. Anesthesiology 106, 312–322

41. Reiner, D. J., Townsend, E. A., Orihuel, J., Applebey, S. V., Claypool, S. M., Banks, M. L., Shaham, Y., and Negus, S. S. (2021) Lack of effect of different pain-related manipulations on opioid self-administration, reinstatement of opioid seeking, and opioid choice in rats. Psychopharmacology (Berl) 238, 1885–1897

42. Gutierrez, T., Oliva, I., Crystal, J. D., and Hohmann, A. G. (2021) Peripheral nerve injury promotes morphine-seeking behavior in rats during extinction. Exp Neurol 338, 113601

43. Guida, F., Iannotta, M., Misso, G., Ricciardi, F., Boccella, S., Tirino, V., Falco, M., Desiderio, V., Infantino, R., Pieretti, G., de Novellis, V., Papaccio, G., Luongo, L., Caraglia, M., and Maione, S. (2022) Long-term neuropathic pain behaviors correlate with synaptic plasticity and limbic circuit alteration: a comparative observational study in mice. Pain 163, 1590–1602

44. Martinez, E., Lin, H. H., Zhou, H., Dale, J., Liu, K., and Wang, J. (2017) Corticostriatal Regulation of Acute Pain. Front Cell Neurosci 11, 146

45. Huang, S., Zhang, Z., Gambeta, E., Xu, S. C., Thomas, C., Godfrey, N., Chen, L., M’Dahoma, S., Borgland, S. L., and Zamponi, G. W. (2020) Dopamine Inputs from the Ventral Tegmental Area into the Medial Prefrontal Cortex Modulate Neuropathic Pain-Associated Behaviors in Mice. Cell Rep 31, 107812

46. Markovic, T., Pedersen, C. E., Massaly, N., Vachez, Y. M., Ruyle, B., Murphy, C. A., Abiraman, K., Shin, J. H., Garcia, J. J., Yoon, H. J., Alvarez, V. A., Bruchas, M. R., Creed, M. C., and Moron, J. A. (2021) Pain induces adaptations in ventral tegmental area dopamine neurons to drive anhedonia-like behavior. Nat Neurosci 24, 1601–1613

47. Mecca, C. M., Chao, D., Yu, G., Feng, Y., Segel, I., Zhang, Z., Rodriguez-Garcia, D. M., Pawela, C. P., Hillard, C. J., Hogan, Q. H., and Pan, B. (2021) Dynamic Change of Endocannabinoid Signaling in the Medial Prefrontal Cortex Controls the Development of Depression After Neuropathic Pain. J Neurosci 41, 7492–7508

48. Bobadilla, A. C., Dereschewitz, E., Vaccaro, L., Heinsbroek, J. A., Scofield, M. D., and Kalivas, P. W. (2020) Cocaine and sucrose rewards recruit different seeking ensembles in the nucleus accumbens core. Mol Psychiatry 25, 3150–3163

49. Bossert, J. M., Stern, A. L., Theberge, F. R., Cifani, C., Koya, E., Hope, B. T., and Shaham, Y. (2011) Ventral medial prefrontal cortex neuronal ensembles mediate context-induced relapse to heroin. Nat Neurosci 14, 420–422

50. Bobadilla, A. C., Heinsbroek, J. A., Gipson, C. D., Griffin, W. C., Fowler, C. D., Kenny, P. J., and Kalivas, P. W. (2017) Corticostriatal plasticity, neuronal ensembles, and regulation of drug-seeking behavior. Prog Brain Res 235, 93–112

51. Cameron, C. M., and Carelli, R. M. (2012) Cocaine abstinence alters nucleus accumbens firing dynamics during goal-directed behaviors for cocaine and sucrose. Eur J Neurosci 35, 940–951

52. Tunstall, B. J., and Kearns, D. N. (2016) Cocaine can generate a stronger conditioned reinforcer than food despite being a weaker primary reinforcer. Addict Biol 21, 282–293

53. Vierck, C. J., and Yezierski, R. P. (2015) Comparison of operant escape and reflex tests of nociceptive sensitivity. Neurosci Biobehav Rev 51, 223–242

54. Deuis, J. R., Dvorakova, L. S., and Vetter, I. (2017) Methods Used to Evaluate Pain Behaviors in Rodents. Front Mol Neurosci 10, 284

55. Huang, J., Zhang, Z., and Zamponi, G. W. (2020) Pain: Integration of Sensory and Affective Aspects of Pain. Curr Biol 30, R393–R395

56. Gourlay, D. L., Heit, H. A., and Almahrezi, A. (2005) Universal precautions in pain medicine: a rational approach to the treatment of chronic pain. Pain Med 6, 107–112

57. Ballantyne, J. C. (2017) Opioids for the Treatment of Chronic Pain: Mistakes Made, Lessons Learned, and Future Directions. Anesth Analg 125, 1769–1778

58. Friedman, R., Li, V., and Mehrotra, D. (2003) Treating pain patients at risk: evaluation of a screening tool in opioid-treated pain patients with and without addiction. Pain Med 4, 182–185

59. Cheatle, M. D., Gallagher, R. M., and O’Brien, C. P. (2018) Low Risk of Producing an Opioid Use Disorder in Primary Care by Prescribing Opioids to Prescreened Patients with Chronic Noncancer Pain. Pain Med 19, 764–773

